# Single-nucleus RNA Sequencing and Spatial Transcriptomics Reveal the Immunological Microenvironment of Cervical Squamous Cell Carcinoma

**DOI:** 10.1101/2021.12.23.473944

**Authors:** Zhihua Ou, Shitong Lin, Jiaying Qiu, Wencheng Ding, Peidi Ren, Dongsheng Chen, Jiaxuan Wang, Yihan Tong, Di Wu, Ao Chen, Yuan Deng, Mengnan Cheng, Ting Peng, Haorong Lu, Huanming Yang, Jian Wang, Xin Jin, Ding Ma, Xun Xu, Yanzhou Wang, Junhua Li, Peng Wu

**Affiliations:** BGI-Shenzhen, Shenzhen 518083, China; Cancer Biology Research Center (Key Laboratory of the Ministry of Education), Tongji Hospital, Tongji Medical College, Huazhong University of Science and Technology, Wuhan 430000, China; Shenzhen Key Laboratory of Unknown Pathogen Identification, BGI-Shenzhen, Shenzhen 518083, China; Department of Gynecologic Oncology, Tongji Hospital, Tongji Medical College, Huazhong University of Science and Technology, Wuhan 430000, China; College of Life Sciences, University of Chinese Academy of Sciences, Beijing 100049, China; College of Innovation and Experiment, Northwest A&F University, Yangling 712100, China; School of Basic Medicine, Qingdao University, Qingdao 266071, China; Department of Biology, University of Copenhagen, Copenhagen DK-2200, Denmark; Department of Obstetrics and Gynecology, Southwest Hospital, Third Military Medical University, Chongqing 400038, China; China National GeneBank, BGI-Shenzhen, Shenzhen 518120, China; Guangdong Provincial Key Laboratory of Genome Read and Write, Shenzhen 518120, China; James D. Watson Institute of Genome Sciences, Hangzhou 310058, China

**Keywords:** cervical cancer, tumor microenvironment, cancer-associated fibroblasts, spatial transcriptomics, single-nucleus RNA sequencing

## Abstract

Effective treatment of advanced invasive cervical cancer remains challenging nowadays. Herein, single-nucleus RNA sequencing (snRNA-seq) and SpaTial Enhanced REsolution Omics-sequencing (Stereo-seq) technology are used to investigate the immunological microenvironment of cervical squamous cell carcinoma (CSCC), a major type of cervical cancers. The expression levels of most immune checkpoint genes in tumor and inflammation areas of CSCC were not significantly higher than those in the non-cancer samples except for *LGALS9* and *IDO1*. Stronger signals of CD56+ NK cells and immature dendritic cells are found in the hypermetabolic tumor areas, while more eosinophils, immature B cells, and Treg cells are found in the hypometabolic tumor areas. Moreover, a cluster of cancer-associated fibroblasts (CAFs) are identified around some tumors, which highly expressed *ACTA2, POSTN, ITGB4,* and *FAP*. The CAFs might support the growth and metastasis of tumors by inhibiting lymphocyte infiltration and remodeling the tumor extracellular matrix. Furthermore, CAFs are associated with poorer survival probability in CSCC patients and might be present in a small fraction (∼20%) of advanced cancer patients. Collectively, these findings might enhance understanding of the CSCC immunological microenvironment and shed some light on the treatment of advanced CSCC.

## 1. Introduction

Cervical cancer is the fourth most common cancer affecting women’s health globally, especially in low- and middle-income regions.^1,2^ Currently, over 12 types of human papillomaviruses (HPVs) are carcinogenic.^3^ Among them, HPV16 is responsible for 60%-70% of the cervical cancer cases, especially cervical squamous cell carcinoma (CSCC). Since 2018, the World Health Organization (WHO) has called for the global elimination of cervical cancer, quantifying actions in vaccination, screening, and disease treatment/management,^4^ which require joint efforts from different parties for decades.

Although early stage cervical cancer receiving radical hysterectomy can achieve a favorable prognosis, the 5-year overall survival rate or disease-free survival rate of advanced cervical cancer are unsatisfactory.^5,6^ At present, chemotherapy (e.g., paclitaxel, cisplatin, bevacizumab, etc.) and radiotherapy remain the main palliative treatments for metastatic or recurrent patients, with a low response rate (48%) and a short survival period (17 months).^7–10^ Immunotherapy brings new hope to treating incurable cervical cancer by reversing the exhausted or suppressed immune activities. Immune-checkpoint blockade (ICB) drugs targeting programmed cell death 1 (PD1), programmed cell death ligand 1 (PD-L1), and cytotoxic T lymphocyte antigen 4 (CTLA4) are currently under trials for recurrent/metastatic cervical cancers.^11,12^ Unfortunately, the overall response rates to ICB therapy were low, varying from 4% to 26%.^11,13–15^ Clarifying the immune landscape of CSCC, especially the immunosuppression status in TME, may help us better address this phenomenon and adjust our treating strategy for cervical cancers.

Single-cell sequencing and spatial transcriptomics are state-of-the-art tools to unravel the cell heterogeneity and microenvironment of tumors, but applications of such techniques to CSCC investigation remain limited. In this study, we collected cervical samples from 20 individuals and combined single-nucleus RNA sequencing (snRNA-seq) and SpaTial Enhanced REsolution Omics-sequencing (Stereo-seq) technology to investigate the immunological profiles of CSCC.^16^ Deciphering the immunological microenvironment of CSCC would provide new insights into the treatment of advanced CSCC, which may accelerate cervical cancer elimination.

## 2. Results

### 2.1 snRNA-seq data revealed the cellular composition of CSCC

To fully characterize the cell composition of cervical tissues, we collected CSCC samples from 5 patients for snRNA-seq (**Figure 1A** **and S1, Table S1**). A total of 67,003 cells and 30,996 genes passed quality control (**Figure 1B, Table S2**), from which we identified 14 cell types based on canonical cell markers (**Figure 1B, Table S3**), including cancer cells (6,960), columnar epithelial cells (CECs, 22,396), endothelial cells (6,340), smooth muscle cells (4,502), fibroblasts (9,836), B cells (689), monocytes (5,281), T cells (4,930), regulatory T (Treg) cells (1,081), plasma cells (3,236), myeloid dendritic cells (DCs) (955), plasmacytoid DCs (272), mast cells (384) and natural killer (NK) cells (141). The uterine cervix contains two types of cells lining its surface, with stratified squamous epithelial cells on the ectocervix and simple columnar epithelial cells on the endocervix and crypts. The dysplasia of squamous epithelial cells leads to CSCC. Therefore, the cancer cells mainly expressed a known CSCC-associated gene *SERPINB3* (Serpin Family B Member 3),^17,18^ tumor gene *TP63*, *CDKN2A,* and keratin gene *KRT15* of squamous cells (**Figure 1C&D**). Since the tissues were mainly from advanced cancer patients (FIGO Stage IB2-IIIC1; the tissue with no staging information was collected during cervical biopsy before chemotherapy), few normal epithelial squamous cells were isolated. Mapping of the snRNA-seq reads against high-risk HPV reference genomes revealed the presence of viral genes in cancer cells (**Figure 1C, Table S2**). We also identified a big cluster of columnar epithelial cells, which highly expressed *MUC5B* and *WFDC2*. This cell type is mainly located in the endocervix epithelia, but it can also appear at the squamocolumnar junction in the adult uterine cervix and some glands. Smooth muscle cells, fibroblasts, and endothelial cells are the major cell types composing the cervical stroma. The smooth muscle cells highly expressed *MYH11, MYLK*, *ACTG2, COL3A1, and COL1A1*, fibroblasts expressed high levels of *LAMA2* besides *COL3A1 and COL1A1*, while endothelial cells can be distinguished by high expressions of *EMCN, FLT1,* and *EGFLT* (**Figure 1C&D**). Besides the structural cells of the cervix, diverse immune cell types were also identified, with monocytes (*ITGAX, MX4A7*), T cells (*CD3E, CD247*), and plasma cells (*MZB1, IGKC*) being the most abundant (**Figure 1B&C**). In short, the snRNA-seq data revealed the structural and immune cell composition of the CSCC tissues, facilitating our downstream spatial transcriptomic analysis with specific cellular gene expression profiles.

**Figure 1.**
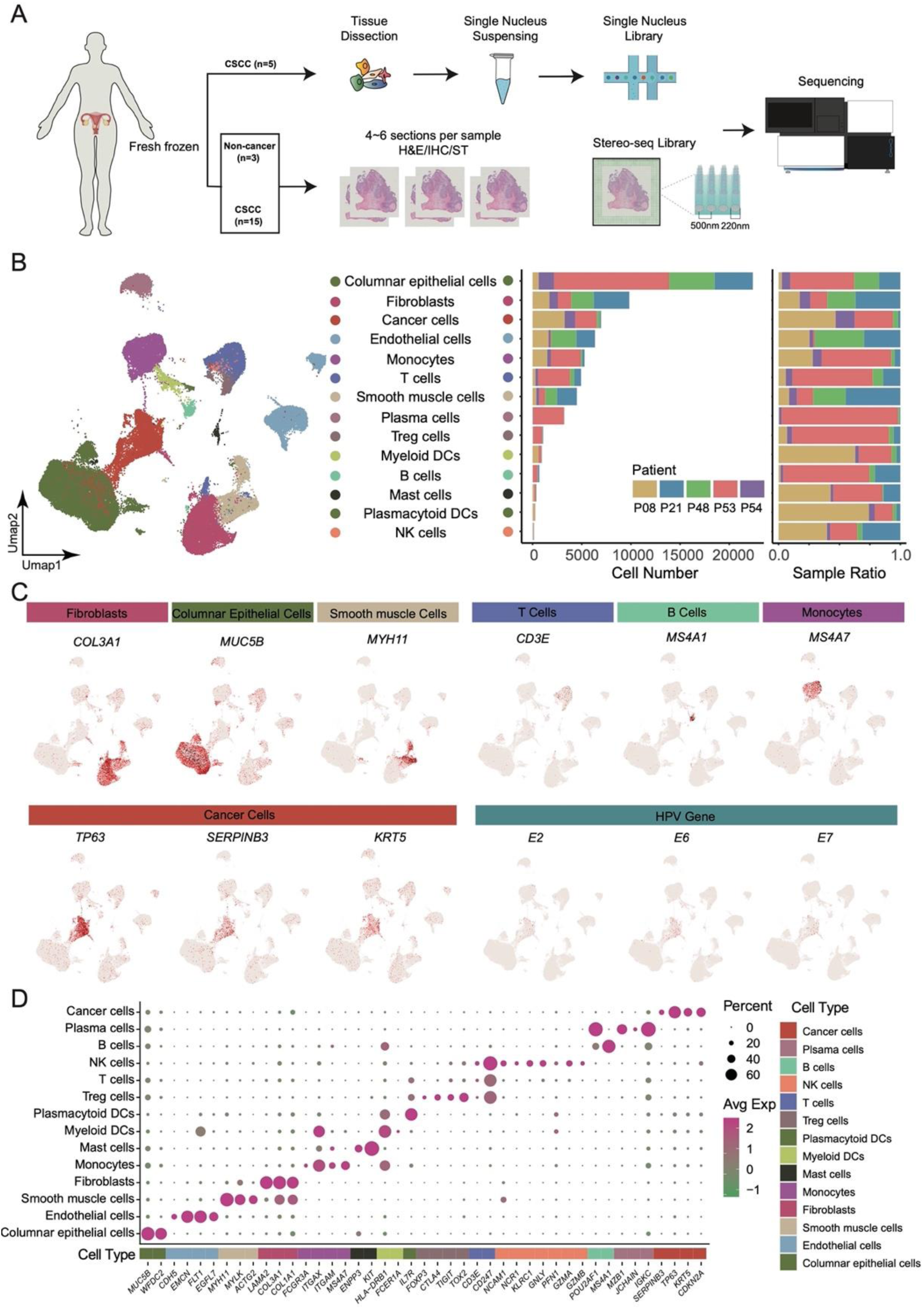
The cellular composition of CSCC tissues. (A) Workflow of snRNA-seq and Stereo-seq experiments applied to cervical tissues. n indicates the number of samples. (B) UMAP of cells identified from the snRNA-seq data of five CSCC tissues (left). The cell number and proportion of each cell type from each sample (right). (C) Expression of selected marker genes and HPV genes in the major cell types of CSCC tissues. (D) Expression matrix of cell-type marker genes in the 14 cell types isolated from CSCC tissues.

### 2.2 Spatial transcriptomic characterization of CSCC

Spatial information is critical in understanding cell-cell interactions in tissues, which unfortunately was missing in the snRNA-seq data. Therefore, we utilized Stereo-seq to acquire the in situ gene expression profiles.^16^ The Stereo-seq chips contained capture probe contained a 25bp coordinate identity barcode, a 10bp molecular identity barcode, and a 22bp polyT tail for mRNA hybridization. Cervical samples from 2 non-cancer and 14 CSCC patients were obtained and embedded in OCT (**Table S1**). Serial cryosections of 10 μm thickness were dissected from each OCT block for Stereo-seq, hematoxylin and eosin (H&E) staining, and immunohistochemical (IHC) staining (**Figure S1A, Table S1**). Finally, a total of 18 Stereo-seq slides were successfully obtained (**Figure 2A, Table S4**). These included 3 slides from 2 non-cancer patients and 15 slides from 14 CSCC patients. Two patients contributed more than one sample. Because the samples were all from different anatomical sites, they were all included to compensate for the small sample size. The capture spots in Stereo-seq chips were 220 nm in diameter with a center-to-center distance of 500 nm between two adjacent spots. The capture spots were grouped into bins to include sufficient genes for accurate clustering. Our preliminary analysis revealed much higher RNA abundance in tumor areas than in stroma areas. To balance the expression differences between tumor and stroma, we annotated the CSCC Stereo-seq slides at bin100 (100 x 100 spots) to fully demonstrate the tissue composition, which would cover an area of approximately 49.72 x 49.72 μm. The mean numbers of genes per bin for the CSCC Stereo-seq slides ranged from 1,767 to 4,152 (**Table S4**). Because the three Stereo-seq slides from non-cancer patients had lower gene expression intensity than CSCC slides, they were annotated at bin200 (99.72 x 99.72 μm). Uniform manifold approximation and projection (UMAP) analysis showed that bin clusters of CSCC and non-cancer tended to dissociate from each other while those of CSCC displayed some convergence (**Figure S1B**).

**Figure 2.**
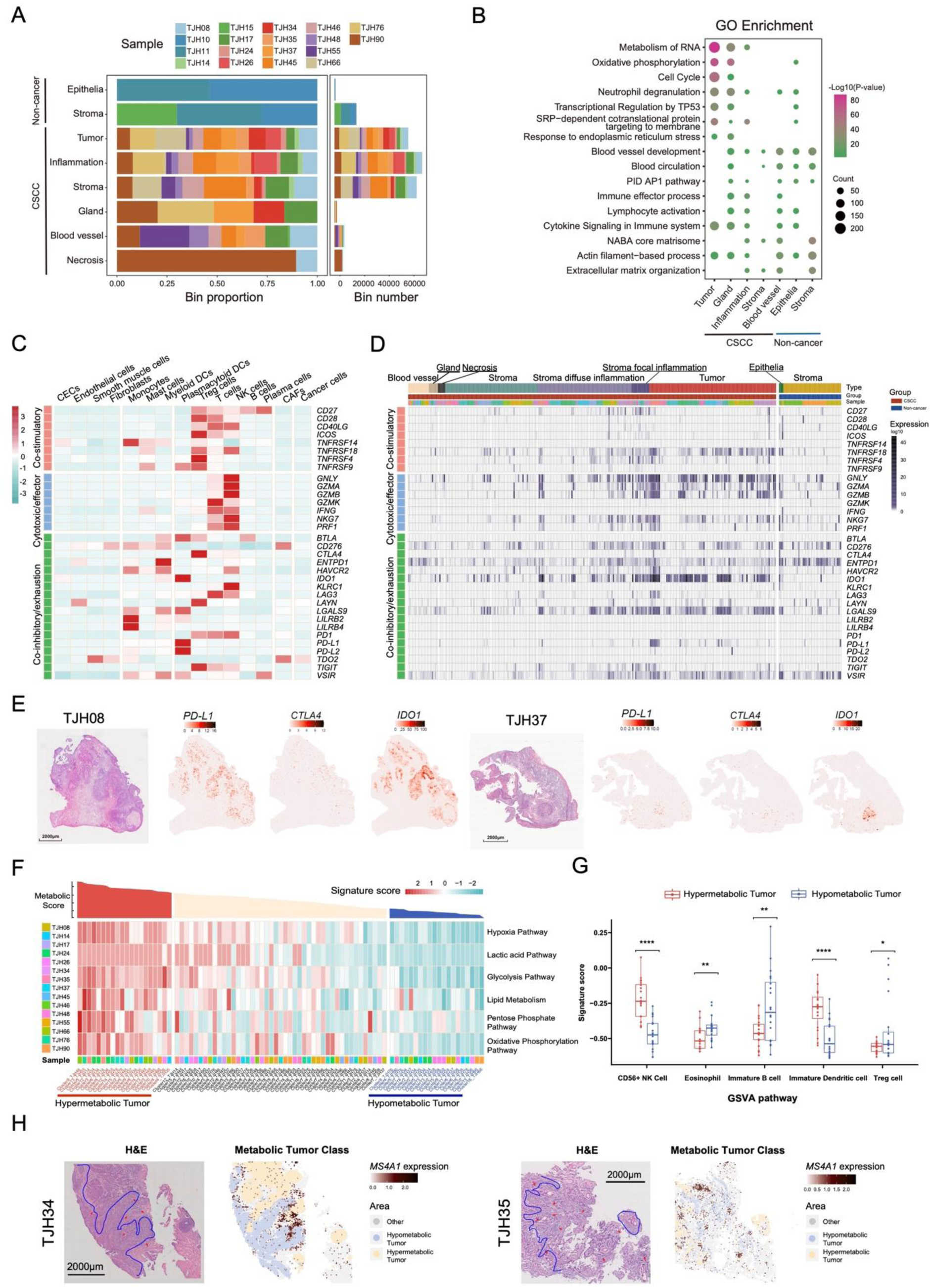
Transcriptomic analysis of the immune heterogeneity in CSCC. (A) Distribution of Stereo-seq clusters in the 18 cervical samples. CSCC (n=15): tumor, stroma, inflammation, gland, blood vessel, and necrosis; non-cancer (n=3, TJH10, TJH11 and TJH15): epithelia and stroma. (B) Gene ontology (GO) enrichment of major Stereo-seq clusters. (C) Heatmap showing the expressions of gene sets associated with different immune functions (co-stimulatory, cytotoxic/effector, and co-inhibitory/exhaustion) in cell types identified by snRNA-seq data. (D) Heatmap showing the expressions of gene sets associated with different immune functions (co-stimulatory, cytotoxic/effector, and co-inhibitory/exhaustion) in Stereo-seq clusters. (E) Spatial expression of selected immune checkpoint genes (*PD-L1, CTLA4,* and *IDO1*) in two representative Stereo-seq samples. The gene tended to be enriched in tumor areas. (F) Top, metabolic scores corresponding to the tumor areas of the 15 CSCC samples. Bottom, heatmap showing the GSVA scores of hypoxia, lactic acid, glycolysis, lipid metabolism, pentose phosphate, and oxidative phosphorylation pathways for tumor areas from each Stereo-seq sample. (G) Violin plots showing abundance of CD56+ NK cells, eosinophils, immature B cells, immature dendritic cells and Treg cells in hyper- and hypo-metabolic tumors. The Y axis shows the GSVA scores for each cell type. The p-values were determined by the Wilcoxon signed-rank test: ns, not significant; *, p <0.05; **, p < 0.01; ***, p < 0.001; ****, p < 0.0001. (H) Spatial expression of marker genes for proliferating cells (*TOP2A*) and B cells (*MS4A1*) in representative two Stereo-seq slides. In the HE images, the blue line indicates the approximate border between hyper- and hypometabolic tumor areas, while the red arrow indicates focal lymphocyte infiltration.

Considering the complex structure of cancerous tissues, we manually conducted the initial annotations of Stereo-seq slides according to professional pathological assessments (based on H&E and IHC staining results) and marker gene expression patterns (**Figure S1C&D)**. Six types of tissue clusters were generally identified in our CSCC samples, including tumor, stroma (without obvious inflammation), inflammation (stroma with diffuse inflammation or focal inflammation), gland, blood vessel, and necrosis. Tumor, stroma, and inflammation clusters were widely distributed among the Stereo-seq slides of the CSCC samples, with certain samples containing necrosis, glands, and blood vessels (**Figure 2A**). Depending on the gene expression profiles, the tissue clusters may be further divided into sub-clusters with number suffixes (**Figure S1C**). It should be kept in mind that a spatial cluster was virtually a mixture of cells in a defined area but was designated by its major characteristics. For example, the tumor clusters were not purely composed of cancer cells, but may also contain other cell types though in small numbers, such as immune cells, fibroblasts, etc. Tissue-specific genes also displayed spatial patterns. Cancerous squamous cell-associated genes such *KRT5, CDKN2A,* and *SERPINB3* were mainly enriched in the Stereo-seq tumor areas, *IGKC* and *IGLC2* enriched in inflammation areas, *VIM* in stromal areas, *ADRA2A* in blood vessels, and *MUC5B* in glands (**Figure S1D and S2**). All the 14 CSCC patients donating the 15 samples were HPV-positive, with 85.7% (12/14) infected by HPV16, 7.1% (1/14) by HPV33, and 7.1% (1/14) by HPV58. The HPV reads covered 8% - 100% (about 600bp - 7905bp) of the viral genome and were mainly identified in the tumor areas (**Table S4, Figure S3 and S4**). Spatial visualization demonstrated varied capture signals of viral genes in different tumor sites of the same sample, with E5, E6, E7, and L1 frequently observed (**Figure S4**). In contrast, only marginal HPV reads were identified in the non-CSCC samples (**Table S4**). In general, higher transcriptional and translational activities, cell proliferation, oxidative phosphorylation, and immune responses were observed in the Stereo-seq tumor areas than in the other areas (**Figure 2B**). The initial manual annotations of the Stereo-seq slides were used to assist downstream analysis of the TME.

### 2.3 Variable immune inhibition in CSCC

Regarding the low response rates of ICB therapy to cervical cancer,^10^ we decided to scrutinize the immune landscape of CSCC for clues. The expression profiles of three gene sets with different immune functions, i.e., co-stimulatory, cytotoxic/effector, and co-inhibitory/exhaustion were evaluated in both snRNA-seq (**Figure 2C**) and Stereo-seq data (**Figure 2D, Table S5**). At the single-cell level, the co-stimulatory genes were expressed in cells of both the innate and adaptive immunity, especially in Treg, T, and NK cells (**Figure 2C**). Treg cells were found to highly express *CD27, CD28, CD40LG, ICOS, TNFRSF18, TNFRSF4,* and *TNFRSF9.* While these genes are necessary for the maturation and normal suppressive function of Treg cells, overexpression of *CD27* in Treg cells may restrain the anti-tumor immune response.^19,20^ Spatially, the co-stimulatory genes tended to be enriched in tumor and inflammation areas (**Figure 2D**). Especially, *TNFRSF18 (*also named glucocorticoid-induced TNF receptor*, GITR)* was commonly expressed in both inflammation and tumor areas. However, its expression was also up-regulated in the epithelia of non-cancer samples. In our snRNA-seq data, this gene was mainly detected in Treg, NK, T cells, and mast cells. Although *TNFRSF18* was associated with the immune suppression by Treg cells in tumors,^21,22^ its high spatial expression level may be jointly contributed by multiple types of immunocytes. The immune cytotoxic/effector genes were mainly expressed by T and NK cells, some of which were commonly up-regulated in the inflammation and tumor regions in the Stereo-seq slides, including *GNLY, GZMA, GZMB,* and *NKG7* (**Figure 2D**). These genes were mainly expressed by NK cells (**Figure 2C**), indicating their important role in the cytotoxic response against CSCC. For the co-inhibitory/exhaustion genes, we failed to detect any prominent expression of *CTLA4* and *PD-1* in our Stereo-seq data (**Figure 2D**), although *CTLA4* can be highly expressed by Treg cells and *PD-1* by Treg, T, and NK cells (**Figure 2C**). *PD-L1*, which was mainly expressed by plasmacytoid DCs, was only overexpressed in the tumor or inflammation areas of a small fraction of Stereo-seq samples. While *CD276, ENTPD1, IDO1, LGALS9, and VSIR* were commonly detected in the Stereo-seq samples, only *IDO1* and *LGALS9* seemed to have higher and wider expressions in the CSCC samples compared to the non-cancer samples (**Figure 2D**). These two genes were both expressed by DCs (**Figure 2C**) and could downregulate cytotoxic T cell activity.^23,24^ Whether IDO1 and LGALS9 could be better targets for ICB therapy against CSCC than CTLA4 and PD-L1 remains to be explored. Moreover, when we zoomed in to check the immune genes in the same Stereo-seq slide, their expressions can vary greatly between different tumor areas. In sample TJH08, the tumor areas commonly expressed high *IDO1*, low *PD-L1,* and very low *CTLA4* (**Figure 2E**). In contrast, only one tumor area in sample TJH37 expressed these genes (**Figure 2E** **and S2**). Collectively, although both our snRNA-seq and Stereo-seq data displayed evidence of immune exhaustion in CSCC patients, the immune microenvironment varied considerably between and within patients.

### 2.4 Metabolic statuses of tumors were associated with different immune responses

Metabolism can modulate the immune microenvironment of tumors, which could be putative intervention targets for cancer therapy.^25,26^ We performed gene set variation analysis (GSVA) separately on six pathways, including hypoxia, lactic acid, glycolysis, lipid metabolism, pentose phosphate, and oxidative phosphorylation pathways. The mean of the GSVA scores for the above six pathways was then calculated as the metabolic score of each tumor area. Based on the GSVA metabolic score, the Stereo-seq tumor clusters ranked top 20 were categorized as hypermetabolic tumors, with those ranked the last 20 as hypometabolic tumors (**Figure 2F**). Generally, the hypermetabolic tumors displayed much higher activities in the oxidative phosphorylation, glycolysis, and lactic acid pathway, indicating active aerobic glycolysis in proliferating cancer cells, i.e., the Warburg effect. Moreover, the hypermetabolic tumors were also accompanied by severe hypoxia and active lipid metabolism, suggesting intense oxidative and nutrient stress in fast-growing tumors. To further explore the relationship between metabolism and immune response, the GSVA signature scores for different immunocytes of each bin within the hyper- and hypometabolic tumor areas were calculated. Based on the average signature score in each tumor area, significant differences were observed for several cell types (**Figure 2G**). CD56+ NK cells and immature dendritic cells showed much stronger signals in the hypermetabolic tumor areas than in the hypometabolic ones, indicating that the hypermetabolic tumors might be more prone to be associated with innate immune response. At the same time, there were more eosinophils, immature B cells, and Treg cells in the hypometabolic tumor areas, which might suggest ineffective adaptive immune response. Two of the Stereo-seq samples, TJH34 and TJH35, contained both hyper- and hypometabolic tumor areas. Using *TOP2A* and *MS4A1* as markers for cancer cells and B lymphocytes, respectively, the spatial association between metabolic and B cell distribution was visualized. Disseminating *MS4A1* expression was detected within and outside the hypometabolic tumor areas, consistent with the lymphocyte distribution pattern in H&E images (**Figure 2H**) and the GSVA signature scores (**Figure 2G**).

### 2.5 Identification of a cluster of cancer-associated fibroblasts (CAFs) in CSCC

When exploring the immune differences between the hyper- and hypo-metabolic tumor areas in sample TJH34, we noticed a unique spatial cluster outside the hypermetabolic tumor regions. This cluster was different from most stromal clusters and looked like a ribbon enclosing the tumor. As this cluster was part of the stroma, we closely scrutinized the fibroblasts in our snRNA-seq data. Luckily, we identified a small set of fibroblasts derived from all five samples (**Figure 3A**), which highly expressed reported marker genes for CAFs, including *ACTA2*, *POSTN, ITGB4,* and *FAP* (**Figure 3B**). The CAFs had a lower stemness score than cancer cells and fibroblasts and were in various cell cycle stages (**Figure 3C**). Function enrichments based on the hallmark gene sets (MSigDB v7.4, https://www.gsea-msigdb.org/gsea/msigdb/) showed that CAFs shared common activities with both fibroblasts and cancer cells (**Figure 3D**). CAFs were involved in similar pathways to fibroblasts including UV response down, angiogenesis, myogenesis, and epithelial-mesenchymal transition. For pathways including the p53 pathway, KRAS signaling down, estrogen response, mitotic spindle, G2/M checkpoint, and E2F targets, CAFs showed similar activities to cancer cells. At the gene level, CAFs not only highly expressed marker genes for fibroblasts, such as the collagen protein family (*COL1A1, COL3A1, COL4A1, COL5A2, COL6A3*, etc.), but also marker genes for malignant squamous cells, such as *KRT4* and *KRT13* of the keratin family (**Figure 3E, Table S6**). Though CAFs in cancers might have different origins, the epithelial characteristics indicated that the origin of the CAFs in our CSCC samples might be associated with EMT.

**Figure 3.**
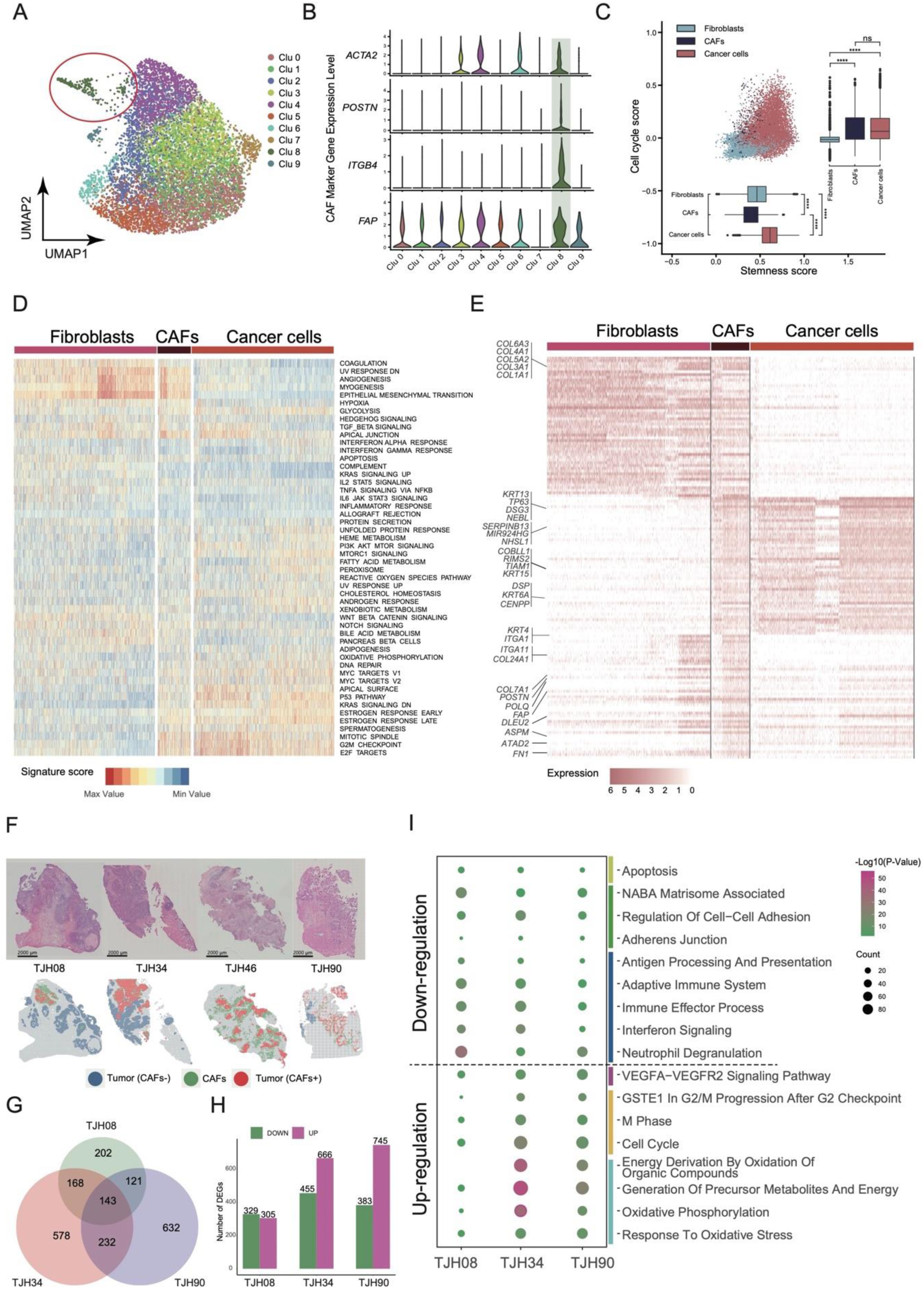
Identification and spatial characterization of cancer-associated fibroblasts (CAFs) in CSCC. (A) UMAP of 9,836 fibroblasts. CAFs were outlined in red. (B) Violin plot showing the expression levels of *ACTA2, POSTN, ITGB4,* and *FAP* in the 10 clusters of fibroblasts. (C) Box plot showing the stemness and cell cycle scores of fibroblasts, CAFs, and cancer cells. The p-values were determined by the Wilcoxon signed-rank test: ns, not significant; *, p <0.05; **, p < 0.01; ***, p < 0.001; ****, p < 0.0001. (D) GSVA results of fibroblasts, CAFs, and cancer cells. (E) DEGs identified in fibroblasts, CAFs, and cancer cells. (F) Spatially projected CAFs in representative Stereo-seq slides. The projected area was determined based on the MIA score of CAFs, *POSTN* expression pattern, and the IHC staining results of POSTN (see **Figure S5B&C**). (G) Venn map showing the number of DEGs identified in the three Stereo-seq samples with both CAFs+ and CAFs-tumors. (H) Bar plot showing the numbers of up- and down-regulated genes in three Stereo-seq samples. The CAFs+ tumors were compared to the CAFs-tumors. (I) Dot plot of enriched GO terms for up- and down-regulated DEGs identified in H.

To locate the spatial distribution of CAFs in CSCC tissues, we adopted the multimodal intersection analysis (MIA) approach developed by Moncada et al. to integrate snRNA-seq and Stereo-seq data.^27^ Briefly, this method calculated the overlapping degree of the expression levels of cell type-specific genes identified by snRNA-seq data and the area-specific genes characterized by Stereo-seq data. The smaller the resultant p-value, which was mentioned as MIA score in our later description, the stronger the correlation between a defined cell type and a spatial area. Initial MIA results showed that our Stereo-seq clustering results complied with the expected cell composition in the corresponding areas (**Figure S5A**). Unfortunately, the MIA score alone cannot fully reflect the spatial specificity of cells, especially in areas with low RNA abundance. Therefore, a high expression level of *POSTN*, which was experimentally verified to be linked with CAFs,^28–30^ and a high MIA score for CAFs were simultaneously utilized to define Stereo-seq clusters of CAFs (**Figure S5B**). Results showed that CAFs were enriched around some tumor areas in 4 out of the 15 Stereo-seq slides (**Figure 3F**), including the hypermetabolic tumor areas of sample TJH34 (**Figure 2H and 3F**). The existence of CAFs in CSCC was further confirmed by IHC staining of POSTN using serial tissue sections of the same samples (**Figure S5C**). Notably, not all the tumor areas were surrounded by CAFs, making us curious about the biological differences associated with the presence of these cells.

### 2.6 CAFs might facilitate the growth and metastasis of CSCC from diverse aspects

To comprehensively reveal the biological functions of CAFs in CSCC, we divided the tumor areas in the Stereo-seq slides into two types: tumor areas surrounded by CAFs (CAFs+ tumors), and tumor areas not surrounded by CAFs (CAFs-tumors). Three Stereo-seq slides were found to contain both CAFs+ and CAFs-tumor areas and were used for downstream analysis. We then used the up- and down-regulated DEGs between the CAFs+ and CAFs-tumor areas (|Log_2_FC| > 1.28, *P <* 0.05) of the 3 samples to perform GO enrichment analysis (**Figure 3G and 3H**, Table S7). Results showed that the CAFs+ tumors were more active in energy usage, metabolism, mitosis, and cell growth than CAFs-tumors (**Figure 3I**). Meanwhile, cellular adhesion, apoptosis, and immune response were down-regulated in CAFs+ tumors. The above observations coincided with the immune and metabolic heterogeneity of CSCC (**Figure 2F&G**), especially in sample TJH34 (**Figure 2H**). These indicated that the presence of CAFs might support tumor progression from different aspects.

Next, we calculated the gene module expression scores of 993 individual bins regarding immune gene sets to evaluate the immune cell abundances. Results showed significantly reduced numbers of B cells, CD4 T cells, CD8 T cells, neutrophils, DCs, NK cells, and Th1 cells in CAFs+ tumors (**Figure 4A** **and S5D**), indicating that CAFs might act as a physical barrier to prevent the infiltration of pro-immunity cells into tumor areas. While more tumor-associated macrophages (TAM) were identified in CAFs+ tumor areas (**Figure 4A**), the distribution of M1 (tumor-suppressive) and M2 (tumor-promoting) phenotypes showed the opposite (**Figure S5E**).^31^ Due to the small sample size and the weak signal of macrophages, we were unsure about the relationship between CAFs and TAMs of different phenotypes.

**Figure 4.**
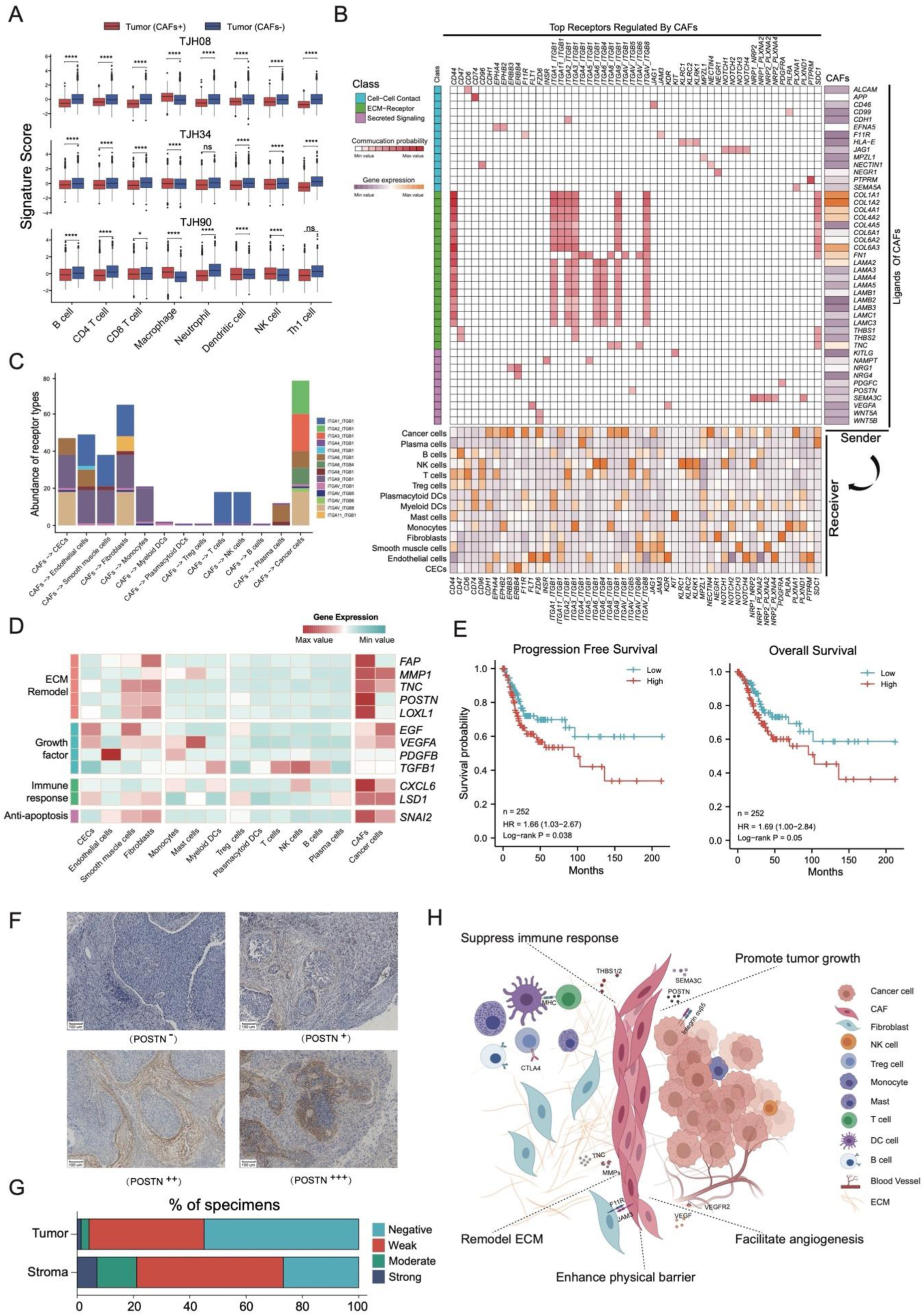
Functional analysis of CAFs. (A) Box plot showing the abundance of immune cells in CAFs+ and CAFs-tumors in three samples. The p-values were determined by the Student’s t-test: ns, not significant; *, p <0.05; **, p < 0.01; ***, p < 0.001; ****, p < 0.0001. (B) Ligand-receptor communication network between CAFs and different cervical cells predicted by snRNA-seq data. The cell–cell communication probability was estimated by integrating gene expression with prior knowledge of the interactions between signaling ligands, receptors and their cofactors. Right, heatmap of the top predicted ligands expressed by CAFs. Middle, heatmap of ligand-receptor pairs between CAFs and different cell types in CSCC. Bottom, expression heatmap of top receptors regulated by CAFs in different cell types. CECs, columnar epithelial cells; ECM, extracellular matrix. (C) Bar plot showing the integrin types involved in potential communications between CAFs and the other cell types. (D) Expression heatmap of gene sets related to functions of CAFs in tumor development in snRNA-seq data. (E) The progression-free and overall survival probabilities of CSCC patients estimated based on the signature scores of the marker gene set for CAFs. The analysis was conducted using the TCGA dataset. (F) Representative IHC staining patterns of POSTN in the stroma adjacent to tumor areas in FFPE CSCC samples. (G) Bar plot showing the IHC staining intensities of POSTN within the tumor area or in the stroma area around tumors in 65 FFPE CSCC samples. (H) Schematic summary of the possible functions of CAFs in CSCC.

As part of the stroma, CAFs will have to interact closely with cancer cells, stromal cells, and immune cells. Indeed, analysis of the snRNA-seq data showed potential interactions between CAFs and the other cells regarding extracellular matrix (ECM) formation and cell-cell contact (**Figure 4B**). CAFs highly expressed genes of the collagen family, especially *COL1A1, COL1A2, COL4A1, COL4A2, COL4A5, COL6A1, COL6A2,* and *COL6A3*) that might interact with *CD44* expressed by immunocytes and smooth muscle cells, which may be involved in cell adhesion and migration. The collagens might also interact with diverse members of the integrin family expressed by the cancer cells, immune cells, and stromal cells. Similarly, *FN1* (fibronectin 1, a soluble glycoprotein) and laminins (*LAMA2, LAMA3, LAMA4, LAMA5, LAMB1, LAMB2, LAMB3, LAMC1, LAMC3*) expressed by CAFs might also interact with the other cell types through integrins. The integrins are membrane receptor proteins made up of α and β subunits and are involved in cell adhesion and recognition. CAFs seems to use different heterodimeric forms of integrins to contact the other cell types. They might interact with cancer cells through integrins composed of subunits α2β1, α3β1, and αvβ8, while interact with endothelial cells, CECs, smooth muscle cells through integrins made up of subunits α9β1, α6β1, and α1β1, and with T cells and NK cells through integrins made up by subunits α1β1 (**Figure 4C**). Importantly, CAFs might take advantage of F11R (also called JAM1, junctional adhesion molecule 1) to form tight junctions with cancer cells and stromal cells through F11R and JAM3, which might prevent the infiltration of immunocytes. CAFs might also express other matrix proteins including *THBS1* (thrombospondin 1), *THBS2,* and *TNC* (tenascin C) to communicate with immunocytes, smooth muscle cells, cancer cells, and plasma cells through *CD44*, integrin (α3β1), and *SDC1*. Moreover, CAFs overexpressed several tissue remodeling factors (**Figure 4D**), including *POSTN* (periostin, a secreted ECM protein), *FAP* (fibroblast activation protein, a serine protease), *MMP1* (matrix metalloproteinase 1), *TNC* (Tenascin-C, a matrix protein), and *LOXL1* (lysyl oxidase like 1, catalyzes the cross-linking of collagen and elastin). This evidence suggested a critical role of CAFs in shaping the tumor extracellular environment.

Besides their role in ECM construction, CAFs might also enhance the stemness and proliferation of cancer cells through overexpressing secreted factors including *SEMA3C, POSTN, CXCL6* (**Figure 4B**). *SEMA3C* was reported to promote cancer stem cell maintenance, angiogenesis, and invasion.^32,33^ *POSTN* can augment cancer cell survival by activating the Akt/PKB pathway through integrins αvβ3.^34^ It may also promote cancer growth through the PTK7-Wnt/β-Catenin signally pathway.^29^ *CXCL6* (C-X-C motif chemokine ligand 6), though mainly related to immune response, was reported to promote the growth and metastasis of esophageal squamous cell carcinoma (**Figure 4D**).^35^ Another highly expressed gene in CAFs, *SNAI2* (Slug), a snail-related zinc-finger transcription factor, may inhibit apoptosis and promote cancer progression.^36,37^ Several common growth factors such as *TGFB1* (transforming growth factor beta 1), *EGF* (epidermal growth factor), and *VEGFA* (vascular endothelial growth factor A) were also expressed by CAFs (**Figure 4D**). Moreover, the upregulation of *LSD1* (histone lysine demethylase 1) in CAFs might inhibit the IFN activation to evade immune attack.^38^ The Wnt5a signaling protein produced by CAFs might also suppress the immune response to facilitate tumor metastasis (**Figure 4B**).^39,40^ In summary, CAFs might be able to potentiate the TME, promoting the progression of tumors.

### 2.7 The presence of CAFs was associated with poorer clinical statuses of CSCC

To verify the pro-tumorigenic effects of CAFs, we first performed survival analyses with a dataset from the Cancer Genome Atlas (TCGA), which contained 252 CSCC patients.^41^ The GSVA score for CAFs was calculated for each CSCC patient using the marker gene set (*ACTA2*, *POSTN, ITGB4,* and *FAP*) (**Figure 3B**). Not surprisingly, higher signals of CAFs predicted unfavorable progression-free survival (HR = 1.66, 95%CI = 1.03-2.67, *P =* 0.038) and overall survival (HR = 1.69, 95%CI= 1.00-2.84, *P =* 0.05) for CSCC patients (**Figure 4E**). Next, we measured the expression levels of POSTN, a biomarker of CAFs, in the stroma and tumor regions using an independent sample set, which consisted of 65 archived formalin-fixed paraffin-embedded (FFPE) CSCC samples (**Figure 4F**). Based on the staining intensity and distribution pattern, we assigned POSTN staining scores to each sample (see Method). The overall positive (weak, moderate, and strong) rate of POSTN expression in the stroma was much higher than that in the tumor regions (**Figure 4G**). However, only 21.1% (14/65) of the FFPE samples had moderate or strong staining of POSTN in the stroma around tumor. Chi-square test showed that higher expression levels of POSTN in the stroma were significantly correlated with more advanced pathological stages, poorer differentiation, larger tumor size, higher squamous cell antigen (SCC) concentrations in peripheral blood, and older age (**Table 1**), which further confirmed the pro-tumorigenic ability of CAFs.

**Table 1.**
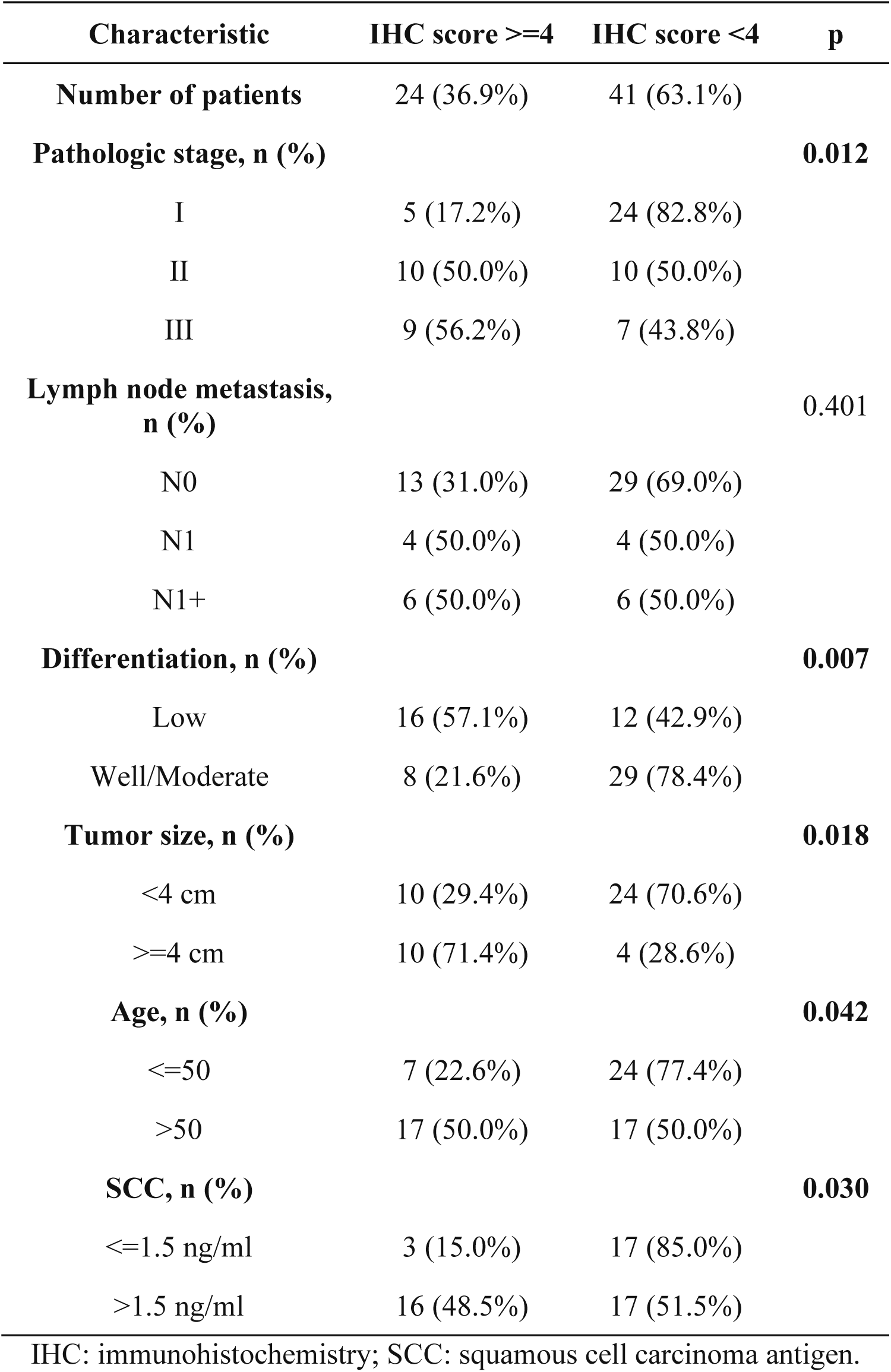
Stromal POSTN expression characteristics in 65 FFPE CSCC samples.

Collectively, our results indicated CAFs were a crucial component of the TME of CSCC, forming a barrier to protect the cancer cells from immune surveillance and clearance. They might also help stimulate cell proliferation and angiogenesis, inhibit apoptosis, and reconstruct the ECM to enhance tumor metastasis (**Figure 4H**).

## 3. Discussion

Although vaccines and radical hysterectomy are effective measures in preventing and treating cervical cancers, the treatment of recurrent/metastatic cervical cancers remains a big obstacle to achieving the goal of cervical cancer elimination. Herein, we have characterized a high-resolution immunological landscape of CSCC combining snRNA-seq and Stereo-seq technology, which may facilitate the management and treatment of HPV-induced cervical cancer.

Nowadays, ICB therapies, especially those using PD-L1/PD-1 and CTLA4 inhibitors, are among the novel methods to treat metastatic cervical cancers. Several studies reported wide expression of PD-L1 in cervical cancers, with positive rates ranging from 34% to 96%.^42–44^ However, PD-L1 expression alone wasn’t associated with the disease outcome of cervical cancer patients.^42^ Indeed, the response rates to PD-1/PD-L1 and CTLA-4 inhibitors fluctuated greatly among different trials and the efficacies seemed independent of the expression status of associated checkpoint genes.^10,45^ In our study, the expression levels of most immune checkpoint genes in tumor and inflammation areas of CSCC were not significantly higher than those in the non-cancer samples except for *LGALS9* and *IDO1* (**Figure 2D**). LGALS9 (i.e., galectin 9) downregulates effector T cell immunity through binding to Tim-3 on the T cell surface or inhibiting the antigen-presenting ability of DCs.^23,24^ While disruption of the galectin 9 signaling pathway was shown to induce tumor regression in mice harboring pancreatic ductal adenocarcinoma, reversed effect was reported in lung metastasis mouse models.^46,47^ IDO1 is mainly expressed in DCs and helps degrade tryptophan into kynurenine, which suppresses T cell functions.^48^ It is found that inhibition of IDO1 could enhance the radiosensitivity of HeLa and SiHa tumorsphere cells, indicating the potential application of IDO1 inhibitors as radiosensitizers.^49^ Whether targeting LGALS9 and IDO1 can improve the treatment of CSCC requires further exploration. Our study also revealed suppressive adaptive immunity in tumor areas with low metabolic activity, highlighting the critical role of metabolic modulation in TME. Indeed, recent clinical trials have combined checkpoint inhibitors with metabolic agents targeting glucose, amino acid, and nucleotide metabolism.^50^ A better understanding of the crosstalk between immune response and metabolism would further benefit cancer therapies.

CAFs have been characterized in multiple types of cancers and can be classified into diverse subtypes.^51^ While cell line studies have implicated a supportive role of CAFs in the proliferation of cervical cancer cells,^52,53^ this is the first study to systemically describe the spatial distribution and biological properties of pro-tumorigenic CAFs in clinical samples of CSCC. Due to tissue heterogeneity, the marker genes for CAFs varied among cancers. In this study, *ACTA2*, *POSTN, ITGB4,* and *FAP* were adequate marker genes to identify CAFs in CSCC (**Figure 4B**). Other genes such as *KRT4, ITGA1, COL24A1,* and *COL7A1* might serve as complementary marker genes for CAFs in CSCC (**Figure 4E, Table S7**), which displayed cellular properties of both fibroblasts and cancerous squamous cells. These CAFs contributed significantly to the heterogeneity of TME, which displayed a pro-tumorigenic phenotype by facilitating tumor growth, metastasis, and immune evasion. The CAFs+ tumors were active in proliferation but lacked lymphocyte infiltration. Exposing these immune-evasive tumors to the immune system is essential to eradicate the cancer cells. However, not all the CSCC samples were positive of CAFs. Only 28.6% (4/14) of the CSCC patients in the Stereo-seq experiment showed presence of CAFs, and only 21.1% (14/65) of the FFPE samples in the IHC experiment were positive of POSTN, a biomarker of CAFs. As CAFs tended to be associated with more advanced pathological status (**Table 1**), chances are that they might function in invasive cancer patients with high variation among individuals. Researchers have tried to interfere with the activation, the action, and the normalization processes of CAFs using antibodies or inhibitors, with several clinical trials ongoing.^54^ Since genes highly expressed by CAFs are also essential to normal tissues, their efficacies and side effects require close monitoring.

Several limitations exist in this study. 1) Samples for snRNA-seq and Stereo-seq were not paired. We were only able to collect paired samples from one cancer patient. Individual and anatomical heterogeneity may hinder the comprehensive annotation of cell types. For example, CAFs were found in 4 Stereo-seq samples, but the number of CAFs was very small in snRNA-seq data. 2) Since all the cervical cancer cases have progressed to invasive stages, most of the tissues collected mainly contained invasive tumors, making it impossible to compare the intra-individual difference between normal epithelia and tumors. 3) Unlike tumor cells, the gene expression levels of immunocytes were relatively low, hindering the spatial analysis of most immune cells at high resolution. 4) Our results were drawn from observations of a limited number of clinical samples. Although we have tried to incorporate public data to validate the pro-tumorigenic phenotype of CAFs, further experimental conformation is necessary.

In conclusion, our data have demonstrated the high heterogeneity of viral gene expression, immune response, and metabolism in CSCC, indicating that combined drugs or therapies targeting multiple biological processes would be better practice to treat CSCC. Interventions on CAFs and tumor metabolism may complement the current treatments of CSCC. Further investigations into these biological aspects may facilitate the development of new drugs or therapies against CSCC and the other HPV-induced squamous cell carcinomas.

## 4. Experimental Section/Methods

### 4.1 Patients and samples

The cervical specimens were collected from 20 patients aged 38 to 69 by the Department of Obstetrics and Gynecology of Tongji Hospital in Wuhan and the Department of Obstetrics and Gynecology of Southwest Hospital in Chongqing. Based on colposcopy examination, 18 patients were diagnosed with CSCC (Stage IB1 to Stage IIIC1), 2 patients were diagnosed with benign gynecological diseases but also required surgery (**Table S1, Figure S1A**). Carcinoma staging was conducted based on the criteria of the FIGO staging system. The freshly-taken samples were used for single-cell RNA sequencing and Stereo-seq experiments. A total of 65 archived FFPE samples were retrospectively obtained from the Department of Obstetrics and Gynecology of Tongji Hospital in Wuhan to verify the presence of CAFs in CSCC.

### 4.2 Experiments

#### 4.2.1 Single-nucleus RNA sequencing

The collected cervical tissues were quick-frozen with liquid nitrogen for 30 minutes and then stored in a -80°C refrigerator. Nuclei isolation and permeabilization were performed under the guidance of Chromium Next GEM Single Cell Multiome ATAC + Gene Expression User Guide (CG000338). snRNA-seq libraries were prepared using the Chromium Single Cell 3ʹ Reagent Kits v3 (10x Genomics, USA), according to the manufacturer’s instructions. Briefly, high-quality sequencing data was obtained after a series of experimental procedures including cell counting and quality control, gel beads-in-emulsion (GEMs) generation and barcoding, post GEM-RT cleanup, cDNA amplification, gene expression library construction, and NovaSeq platform (Illumina, USA) sequencing.

#### 4.2.2 Tissue preparation for spatial transcriptomic experiment

A tissue block with an edge length of less than 1cm was dissected from the surgically removed tissues. The tissue block was then rinsed by cold PBS, immersed in the pre-cooled tissue storage solution (Miltenyi Biotec, Germany), and then embedded with pre-cooled OCT (Sakura, USA) in a -30°C microtome (Thermo Fisher, USA) within 30 minutes after surgery. Three to four serial cryosections of 10 µm thickness were cut from the OCT-embedded samples for H&E staining, Stereo-seq library preparation, and IHC staining. Brightfield images of the H&E samples were taken with a Motic microscope scanner (Motic, China) for histopathological assessment.

#### 4.2.3 Quality control of RNA obtained from OCT-embedded samples

100-200 μm thick sections were cut from each OCT-embedded sample for total RNA extraction using the RNeasy Mini Kit (Qiagen, USA) according to the manufacturer’s protocol. RNA integrity number (RIN) was determined by a 2100 Bioanalyzer (Agilent, USA). Only samples with RIN≥7 were qualified for the transcriptomic study. All samples used had RIN of 7-10.

#### 4.2.4 Stereo-seq library preparation and sequencing

The spatial transcriptomic RNA library was constructed using Stereo-seq capture chips (BGI-Shenzhen, China), which had a size of 1 cm^2^. The capture spots were 220 nm in diameter with a center-to-center distance of 500 nm between each other. Each Stereo-seq capture probe contained a 25bp coordinate identity barcode, a 10bp molecular identity barcode, and a 22bp polyT tail for in situ mRNA hybridization.^16^ A cryosection of 10 μm thickness cut from OCT-embedded tissue was quickly placed on the chip, incubated at 37°C for 3 minutes, and then fixed in pre-cooled methanol at -20°C for 40 minutes. The fixed tissue section was stained with the Qubit ssDNA dye (Thermo Fisher, USA) to check tissue integrity before fluorescent imaging. After that, the tissue section was permeabilized using 0.1% pepsin (Sigma, USA) in 0.01 N HCl buffer, incubated at 37°C for 14 minutes, and then washed with 0.1x SSC. RNA released from the permeabilized tissue was reverse transcribed for 1 hour at 42°C. Later, the tissue section was digested with tissue removal buffer at 42°C for 30 min. The cDNA-containing chip was then subjected to cDNA release enzyme treatment overnight at 55°C. The released cDNA was further amplified with cDNA HIFI PCR mix (MGI, China). Around 20ng cDNA was fragmented to 400-600bp, amplified for 13 cycles, and purified to generate DNA nanoball library, which was sequenced with the single-end 50+100bp strategy on an MGI DNBSEQ sequencer (MGI, China).

#### 4.2.5 Immunohistochemical (IHC) staining

The IHC staining for Ki67 and POSTN were performed under the manufactures’ protocol. The frozen sections dried at room temperature were placed in a 37°C oven for 10-20 minutes, fixed with 4% paraformaldehyde for 20 minutes, and washed thrice with PBS (pH = 7.4) for 5 minutes. The antigens were then repaired with EDTA (pH9.0) and the endogenous peroxidase was blocked by 3% hydrogen peroxide. The slides were further blocked with 3% BSA (G5001-100g, Servicebio) at room temperature for 30 minutes, and then incubated with Ki67 (ab16667, Abcam, 1:200) or POSTN (ab215199, Abcam, 1:500) at 4°C overnight. Finally, the frozen slices were subjected to secondary antibody blocking, DAB staining, nuclear re-staining, and dehydration. The protein expression levels of Ki67 and POSTN were evaluated by professional pathologists under a microscope. The expression scores of POSTN in tumor and stromal regions of 65 FFPE samples were measured according to the positivity percentage (0-5% = 0, 5-25% = 1, 26-50% = 2, 51-75% = 3, >75% = 4) and staining intensity (negative = 0, weak = 1, moderate = 2, strong =3) (**Figure 4F and 4G**). The final score was obtained by multiplying the two scores, ranging from 0 to 12. Negative =0, weak = 1-4, moderate = 4-8, strong = 9-12 (**Table 1**).

### 4.3 Bioinformatic analysis

#### 4.3.1 Cell type characterization using snRNA-seq data

##### Quality control and gene expression quantification of snRNA-seq data

Raw sequencing files were first processed using CellRanger version v6.0.2 (10x Genomics, USA) to obtain gene expression matrices. After cell calling, the droplets containing no cell were excluded based on the number of filtered unique molecular identifiers (UMIs) mapped to each cell barcode. Droplets with low-quality cells or more than one cell were also removed. To obtain a gene expression matrix optimized to individual samples, the R package scCancer v2.2.1 was employed to further filter the expression matrix.^55^ The filtering thresholds were determined by catching outliers from the distribution of four quality spectra, including the number of total UMIs, the number of expressed genes, the percentages of UMIs from mitochondrial genes, and the percentages of UMIs from ribosomal genes. Besides filtering cells, genes expressed in less than three cells were also excluded to avoid false-positive results. The filtering thresholds for the five samples were documented in **Table S2**.

##### Cell type clustering using multi-sample snRNA-seq data

Integrative analysis of the snRNA-seq data from the five patients was carried out using the IntegrateData function of Seurat v4.^56^ Further analysis included normalization, log-transformation, highly variable genes identification, dimension reduction, clustering, and differential expression analysis were all conducted using default parameters of Seurat except that *dims* was set as 1:30. Initially, a total of 35 cell clusters were obtained (69,312 cells with 30,996 genes). To ensure reliable identification, we removed the cell clusters consisting of less than two samples and with less than 15 cells per sample. Finally, 14 cell clusters (67,003 cells with 30,996 genes) were determined based on reported cell type marker genes (**Table S3**).

##### Analysis of differentially expressed genes (DEGs)

Expression of each gene in each cluster was compared against the rest of the clusters using the Wilcoxon rank-sum test with the FindAllMarkers function of Seurat v4.^56^ Significantly up- or down-regulated genes were identified using the following criteria: 1) the difference in gene expression level was >1.28 fold unless explicitly noted; 2) genes were expressed by more than 25% of the cells belonging to the target cluster. 3) the adjusted p-value was less than 0.05.

#### 4.3.2 Processing and annotation of Stereo-seq data

##### Preliminary processing of Stereo-seq data

Stereo-seq raw data were automatically processed using the BGI Stereomics analytical pipeline (http://stereomap.cngb.org/), where the reads were decoded, trimmed, deduplicated, and mapped against the human and HPV reference genomes. The reference genomes were: Human, GRCh38.p12; HPV16:K02718.1; HPV18: EF202147.1; HPV33: M12732.1; HPV58: D90400.1. Data of the chip area covered by tissue was extracted based on the ssDNA and H&E staining images using the Lasso function of the BGI Stereomics website. It’s worth noting that tumor sites usually had much higher overall mRNA levels than the other anatomical areas, leading to a significant imbalance of transcriptomic signals between the tumor areas and the other sites on the Stereo-seq slides. Therefore, to fully reflect the spatial transcriptomic landscape around the tumor areas, a bin size of 100 (100 spots x 100 spots, i.e., 49.72 x 49.72 μm) was used as the analytical unit for the annotation of CSCC Stereo-seq slides, while a bin size of 200 (200 spots x 200 spots, i.e., 99.72 x 99.72 μm) was used for the non-CSCC samples. The downloaded data was then processed with Seurat v4.^56^ We used the criteria of >200 UMIs per bin to remove bins with low expression signals. The data were then normalized using the SCTransform function. Dimension reduction was conducted with principal PCA. Unsupervised clustering of bins was performed with UMAP. Sequencing and analytical details can be found in **Table S4**.

##### Annotation of bin clusters in Stereo-seq slides

The bin clusters were annotated based on the in situ expression patterns of marker genes combining H&E and IHC staining results. The spatial expression patterns of genes in Stereo-seq slides (**Figure S1 and S2**) were conducted with the SpatialFeaturePlot function of Seurat v4.^56^ The H&E and IHC images were examined by professional pathologists to determine the tissue types. The annotated Stereo-seq areas were confirmed to be consistent with the H&E and IHC assessment and marker gene expression patterns.

##### Identification of viral RNA

The viral reads were mapped against HPV reference genomes with BWA. The genome coverage (covered length/full length of the reference genome) and effective depth (total mapped bases/covered length) for each type were calculated. Only samples with a viral genome coverage of >5% and an effective depth of >50x were deemed as HPV positive (**Table S2&S4, Figure S3**).

#### 4.3.3 Characterization of the immunity, hypoxia, and metabolism statuses in CSCC

##### Signature enrichment analysis of Stereo-seq clusters

In enrichment analysis, the expression scores of signature genes were calculated for individual bins using the AddModuleScore (on log-normalized data) of Seurat v4 with default parameters.^56^ Pathways and cell types included in the enrichment analysis, with the corresponding reference for gene signatures, were listed as follows: hypoxia,^57^ glycolysis,^58^ lipid metabolism,^59^ lactic acid metabolism, oxidative phosphorylation (MSigDB, https://www.gsea-msigdb.org/gsea/msigdb/), pentose phosphate pathway,^60^ macrophages,^61^ and other immune cells (**Figure 2F and 2G**).^62^

##### Prediction of the spatial distribution of immunocytes

Using the gene signatures of immunocytes as input, we calculated the cell type scores of each bin in tumor areas using the AddModuleScore of Seurat v4.^56^ Combining with the spatial coordinate of bins, the possible spatial distribution of the corresponding cell type was obtained (**Figure S5D**).

#### 4.3.4 Characterization of CAFs

##### Signature enrichment analysis of fibroblasts, CAFs, and cancer cells with snRNA-seq data

Expression scores of signature genes from MSigDB v7.4 (https://www.gsea-msigdb.org/gsea/msigdb/) were calculated for individual cells using the AddModuleScore function (on log-normalized data) of Seurat v4 with default parameters to assess differential pathways in fibroblasts, CAFs, and cancer cells.^56^

##### Cell cycle and stemness analysis of fibroblasts, CAFs, and cancer cells with snRNA-seq data

We used the AddModuleScore function of Seurat v4 to calculate the relative average expression of a list of G2/M and S phase markers to obtain the cell cycle scores.^63^ Cell stemness analysis was conducted using the OCLR model and the stemness signatures embedded in the scCancer package.^55,64^

##### Multimodal intersection analysis (MIA)

To integrate snRNA-seq and Stereo-seq data, we calculated the overlapping degree of the expression levels of cell type-specific genes identified by snRNA-seq data and the area-specific genes characterized by Stereo-seq data using the MIA approach.^27^ The lower the p-value, the higher overlapping between a certain cell type and a Stereo-seq area. MIA was conducted to confirm the consistency between cell types and Stereo-seq annotated areas and to identify the Stereo-seq areas composed of CAFs.

##### DEG and GO enrichment analysis of CAFs+ and CAFs-tumor areas

Expression of each gene in CAFs+ cluster was compared against that in the CAFs-clusters of the same Stereo-seq chip using the Wilcoxon rank-sum test with the FindMarkers function of Seurat v4.^56^ Significantly up- or down-regulated genes (**Figure 3G**) were identified using the following criteria: 1) the difference in gene expression level was >1.28 fold unless explicitly noted; 2) genes were expressed by more than 25% of the bins belonging to the target cluster. 3) the adjusted p-value was less than 0.05. GO enrichment analysis was conducted using Metascape with default settings (**Figure 3I**).^65^

##### Cell-cell communication between CAFs and the other cell types in CSCC tissues

To understand the communication network between CAFs and the other cell types, we conducted cell-cell communication with CellChat with the snRNA-seq data to obtain the ligand-receptor pairs regulated by CAFs.^66^ The probability of cell–cell communication was estimated by integrating gene expression with prior knowledge of the interactions between signaling ligands, receptors and their cofactors.

##### Prognostic analysis of CAFs with TCGA data

The gene expression profiles of CSCC were downloaded from The Cancer Genome Atlas (TCGA) (https://portal.gdc.cancer.gov/) with the latest follow-up prognostic information obtained from an integrated clinical data resource. We calculated a signature score of CAFs for each CSCC patient with GSVA using the marker genes of CAFs (*ACTA2*, *POSTN, ITGB4,* and *FAP*). Based on the median GSVA score, the patients were then divided into two groups: high CAFs v.s. low CAFs. The Kaplan-Meier overall survival and progression-free survival curves were generated with GraphPad Prism 6 (**Figure 4E**).

### 4.4 Statistical analysis and plotting

Statistical analyses, including Student’s t-test, Wilcoxon’s rank-sum test, Wilcoxon signed-rank test, and Chi-square test were performed in R 3.6.0. Asterisks indicate the significance levels of p-values: ns, not significant; *, p <0.05; **, p < 0.01; ***, p < 0.001; ****, p < 0.0001. The schematic plots of CAFs were created with BioRender (https://biorender.com/).

## Ethical statement

This study was reviewed and approved by the Medical Ethics Committee of Tongji Medical College, Huazhong University of Science and Technology (TJ-IRB20210609), Southwest Hospital, Third Military Medical University (KY2020142), and the Institutional Review Board of Beijing Genomics Institute, Shenzhen, China (BGI-IRB 21050).

## Data and code availability

The data and scripts supporting the findings of this study have been deposited into CNSA (CNGB Sequence Archive) of CNGBdb (https://db.cngb.org/cnsa/, accession numbers to be updated) and are available upon reasonable request. The dataset used to verify the distinguishing ability of the DEGs between preinvasive and invasive cancerous lesions was obtained from the Gene Expression Omnibus (GEO) database (www.ncbi.nlm.nih.gov/geo) by the Accession Number of GSE63514.

## Author contributions

A. Z. Ou, P. Wu, J. Li, W. Ding, and D. Chen designed this study. P. Wu, Z. Ou, J. Li, X. Xu, D. Ma, X. Jin, H. Yang, and J. Wang supervised this study. P. Wu, Y. Wang, W. Ding, and S. Lin coordinated sample collection. S. Lin, Y. Ding, and T. Peng conducted snRNA-seq and IHC experiments. P. Ren, Y. Tong, and D. Wu performed Stereo-seq library construction and H&E staining. A. Chen and M. Cheng provided technical support for Stereo-seq experiments. H. Lu conducted sequencing for Stereo-seq libraries. J. Qiu, J. Wang, Y. Tong, and D. Wu conducted data analysis. Z. Ou, S. Lin, P. Ren, J. Qiu, and J. Wang wrote the original manuscript. Z. Ou, S. Lin, P. Wu, J. Li, W. Ding, and D. Chen reviewed and polished the manuscript.

## Acknowledgements

This work received funding from the National Key R&D Program of China (2021YFC2701201), the National Natural Science Foundation of China (82072895, 82141106, 82103134) and the Guangdong Provincial Key Laboratory of Genome Read and Write (2017B030301011). We thank China National GeneBank for providing sequencing services for this project. The authors also would like to thank Miss Feiyun Xuanyuan and Mr. Geer Xuanyuan for their inspirational communications.

## Supplementary Tables (see Excel files)

**Table S1.** Clinical characteristics and experimental details for samples.

**Table S2.** snRNA-seq data statistics.

**Table S3.** Marker genes for the annotation of cell types in CSCC.

**Table S4.** Stereo-seq data statistics.

**Table S5.** Expression matrix of immune genes in Stereo-seq areas.

**Table S6.** DEGs for fibroblasts, CAFs, and cancer cells.

**Table S7.** DEGs for CAFs+ and CAFs-tumors.

---

**Figure S1.**
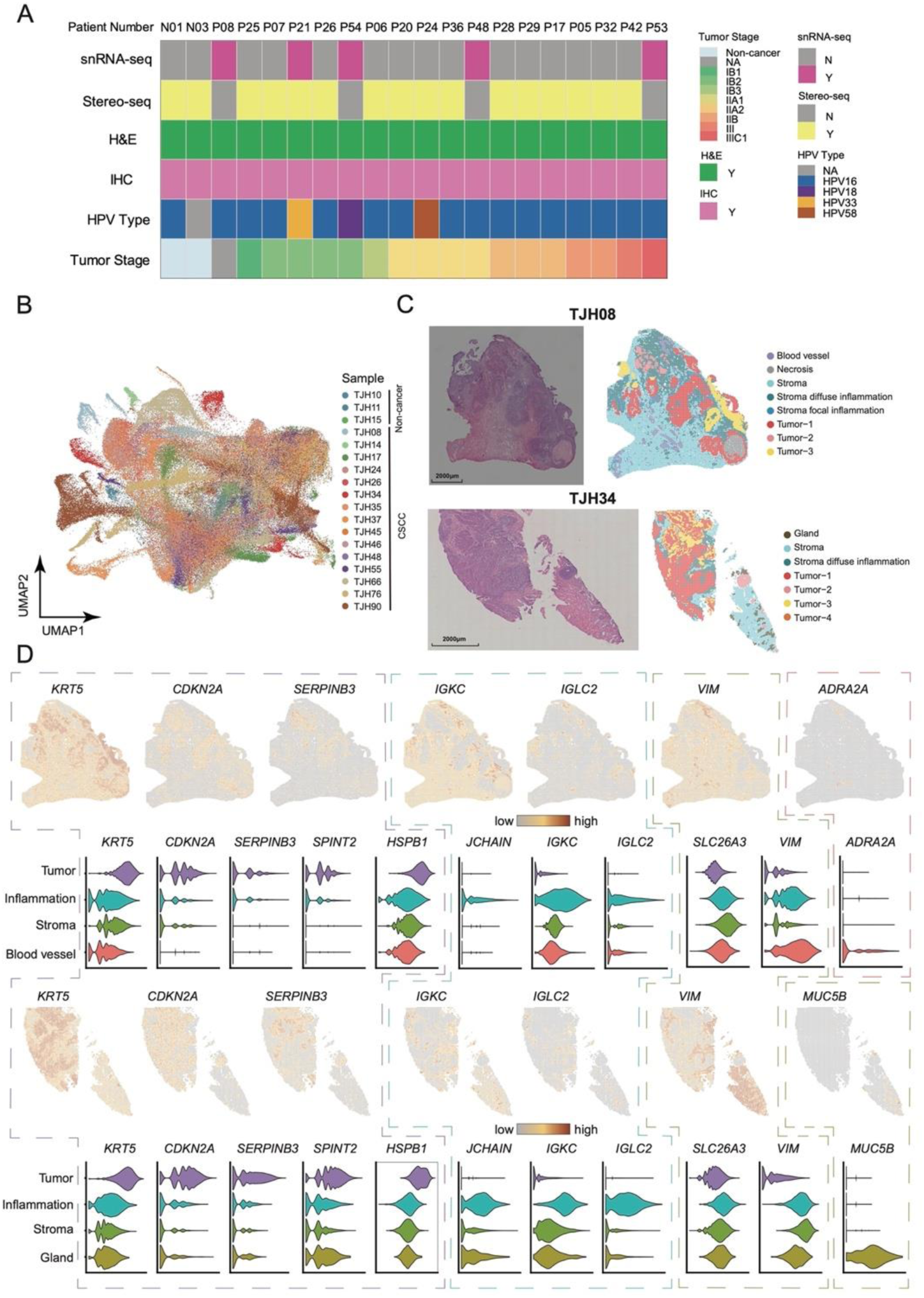
Experimental details and the annotation process of Stereo-seq slides. (A) Clinical characteristics and experimental details of cervical samples from CSCC and non-cancer patients. (B) UMAP of Stereo-seq bins from 18 cervical samples. (C) Annotation results of 2 representative Stereo-seq slides. (D) Expression of tissue-specific genes in 2 representative Stereo-seq slides. The outline color indicates different tissue types: purple, tumor; blue, stroma with inflammation; green, stroma; red, blood vessel; brown, gland. Violin plots display the gene expression levels in the Stereo-seq areas.

**Figure S2.**
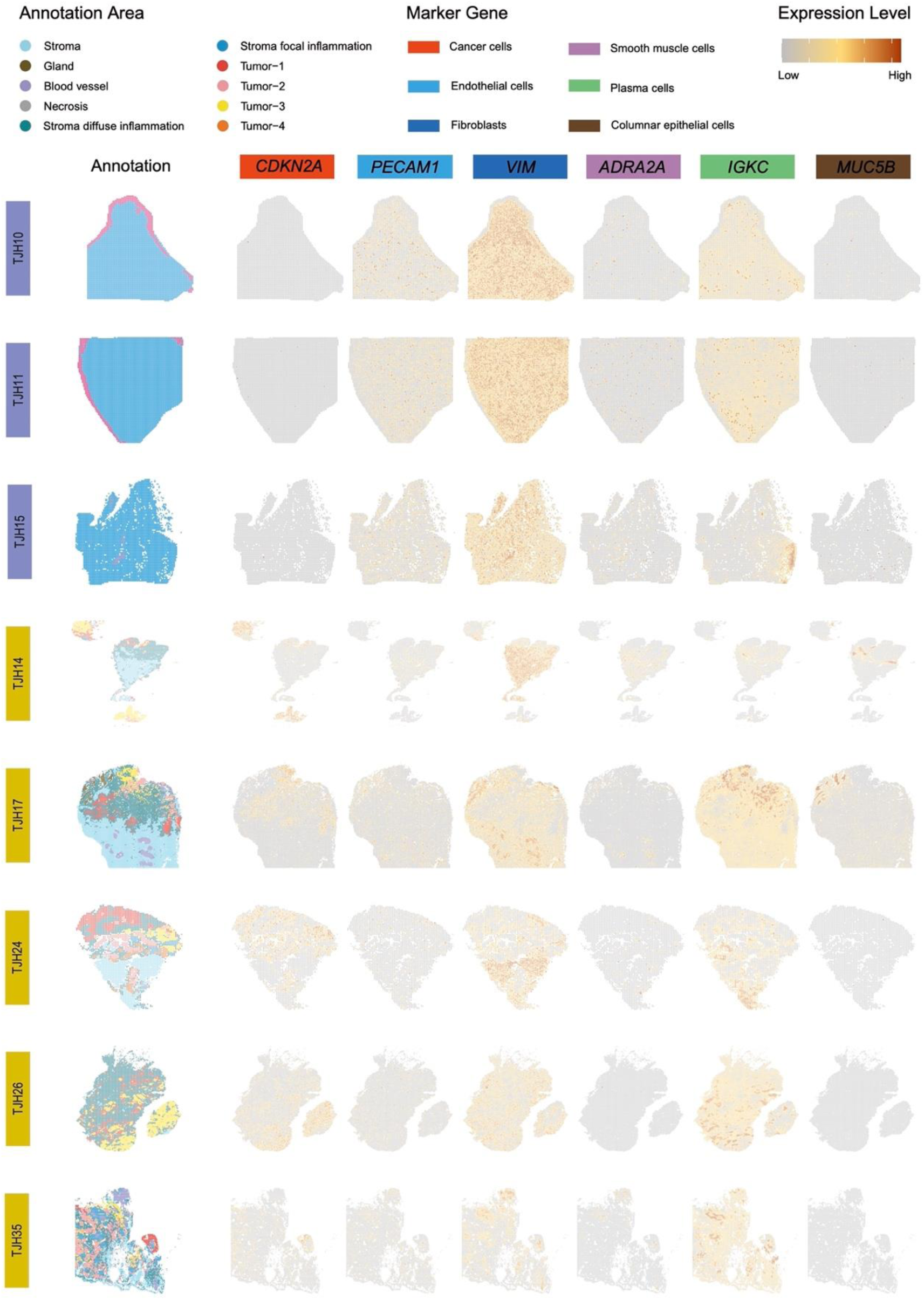
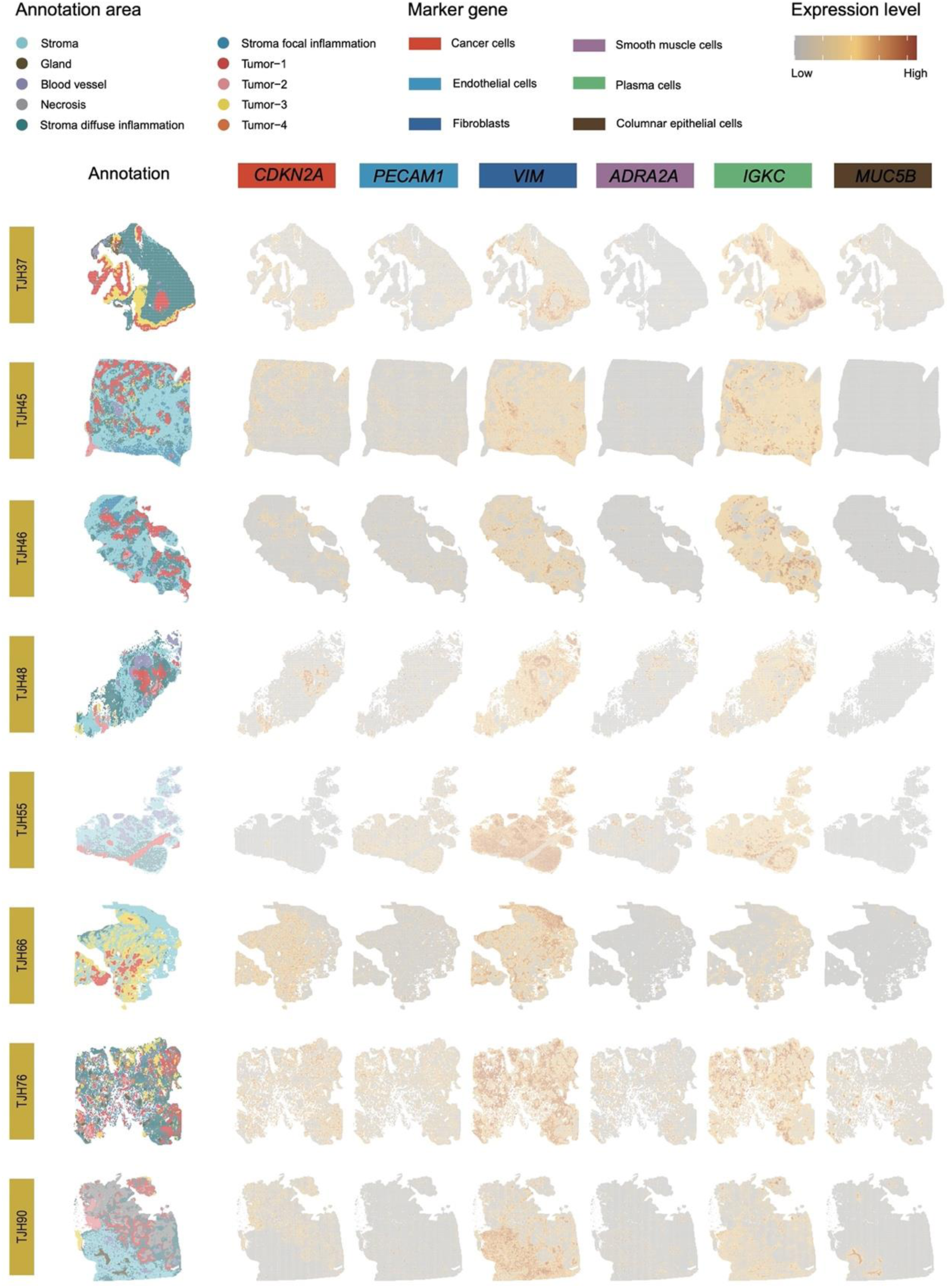
Annotation results of the 18 Stereo-seq slides. Non-cancer samples included TJH10, TJH11, and TJH15. All the other samples were CSCC samples. Six genes were selected to represent different tissue types. Cancer cells, *CDKN2A*; Endothelial cells, *PECAM1*; Fibroblasts, *VIM*; Smooth muscle cells, *ADRA2A*; Plasma cells, *IGKC*; Columnar epithelial cells, *MUC5B*.

**Figure S3.**
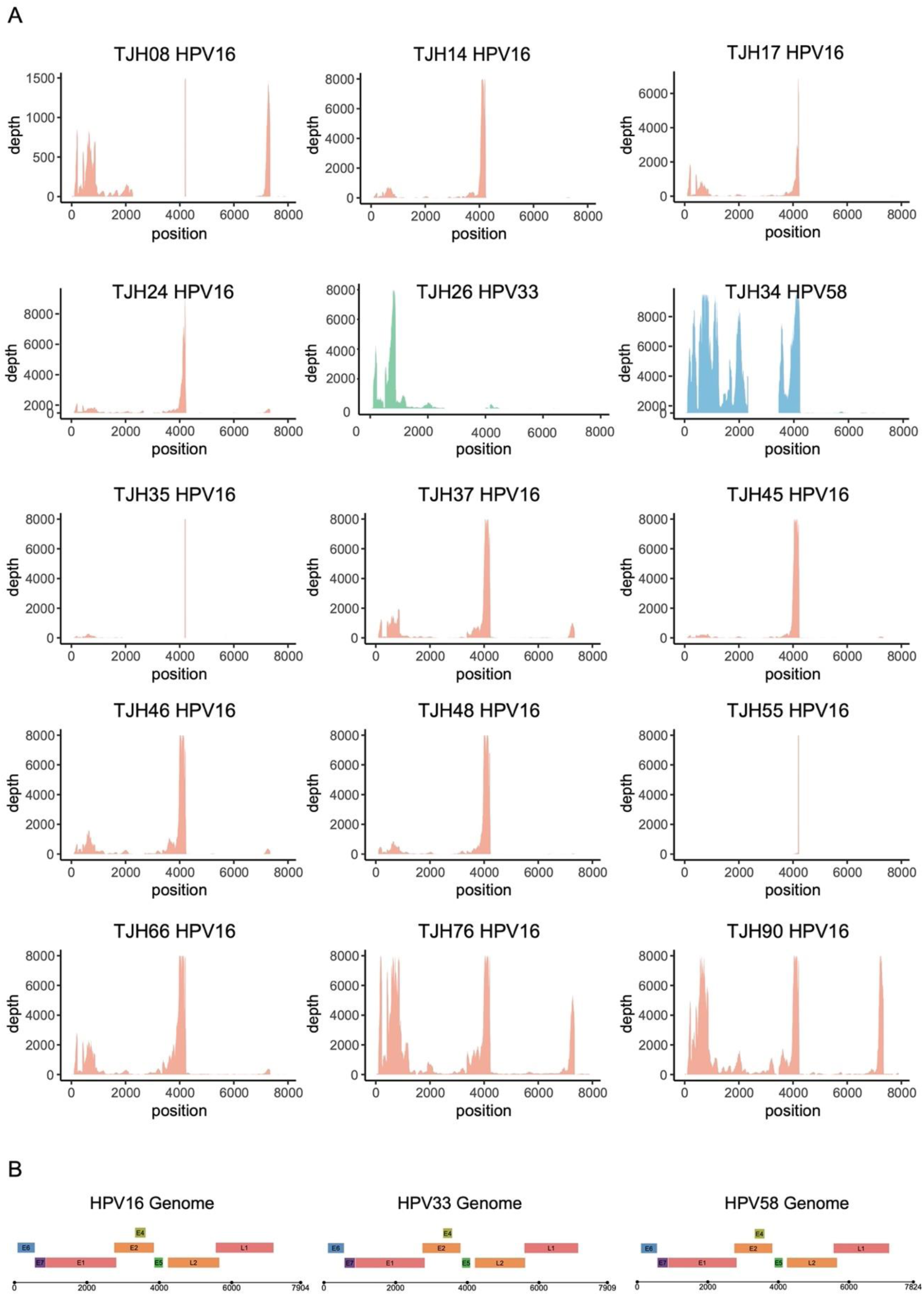
HPV reads in the Stereo-seq sequencing data of 15 CSCC tissues. (A) Mapping of HPV reads against the corresponding reference genome. (B) Schematic plot showing the genomic arrangement of the HPV genes in a linear form.

**Figure S4.**
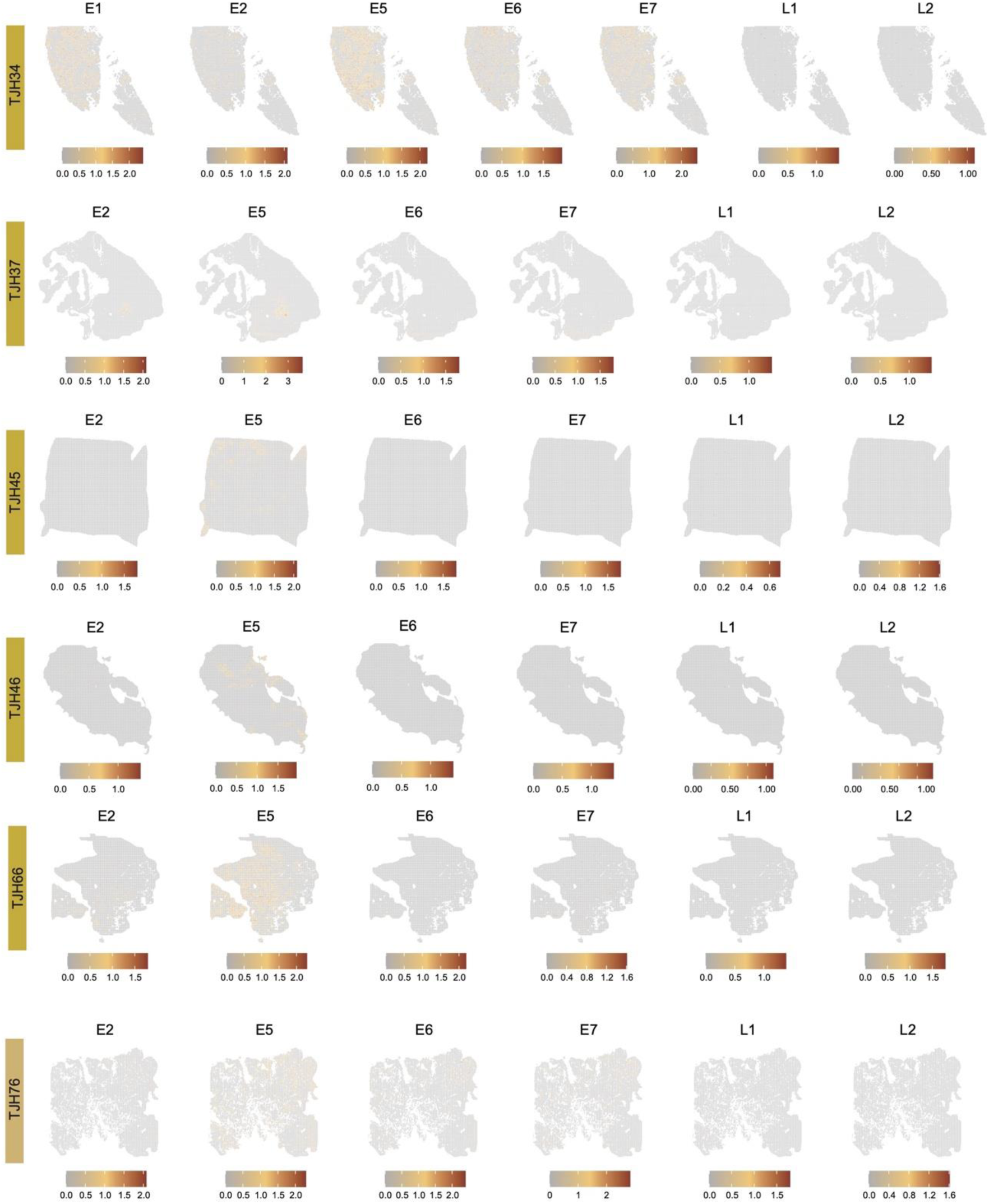
Expression of HPV genes in the tumor areas of selected CSCC Stereo-seq slides. The annotation result for each sample can be found in Figure S2.

**Figure S5.**
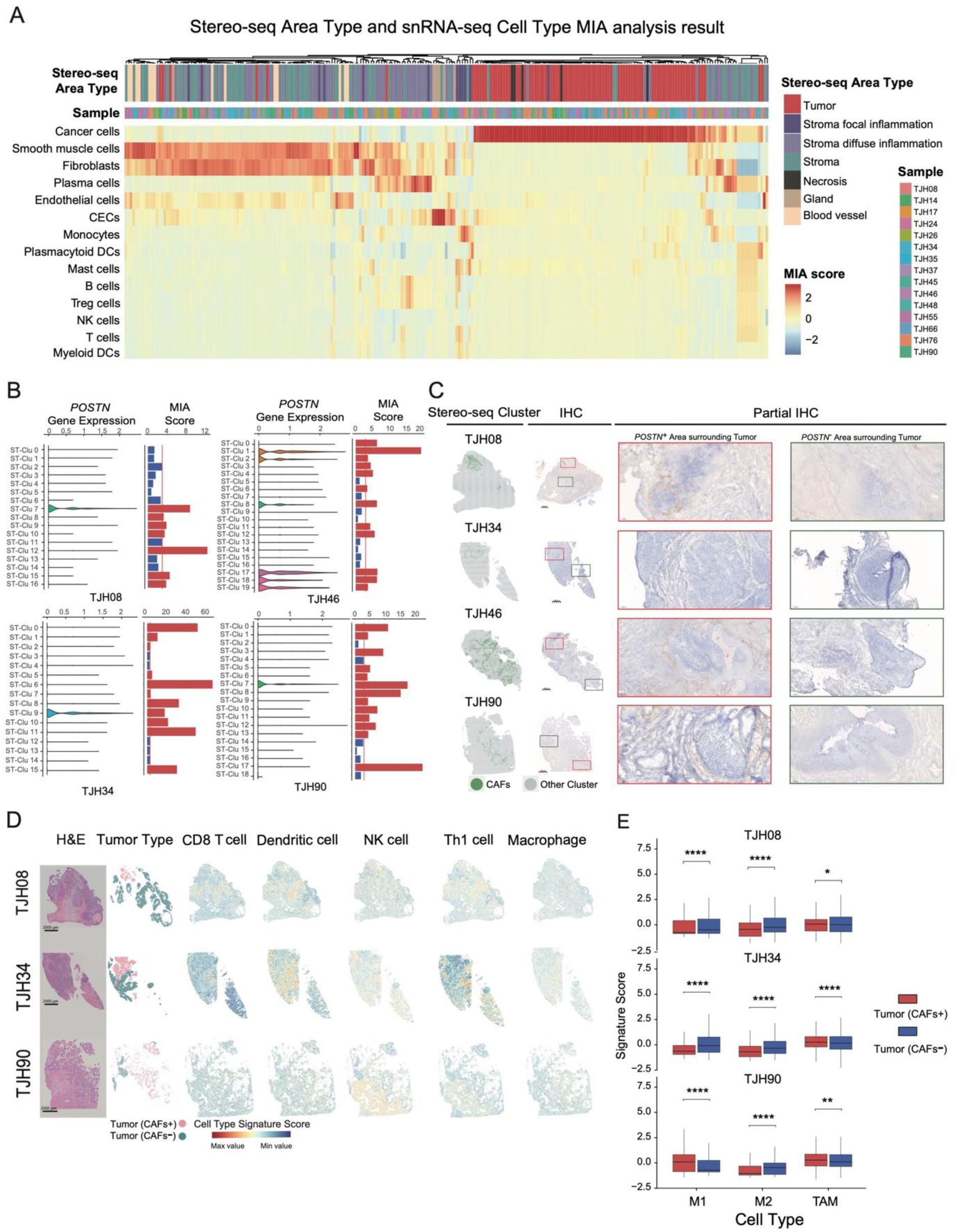
Characterization of CAFs. (A) Correlation between snRNA-seq cell types and the Stereo-seq areas defined by multimodal intersection analysis (MIA). (B) Expression of *POSTN* in Stereo-seq clusters and the associated MIA scores for CAFs. (C) Spatial clustering of CAFs in Stereo-seq slides and the IHC staining of POSTN in corresponding serial sections of CSCC samples. (D) Spatial prediction of immunocytes in Stereo-seq slides. (E) The abundance of macrophages including M1 (tumor-suppressive phenotype macrophage), (tumor-promoting phenotype macrophage) M2, and TAM (tumor-associated macrophage) in CAFs+ and CAFs-tumors. Asterisks indicate the significance levels of p-values: *, p <0.05; **, p < 0.01; ***, p < 0.001; ****, p < 0.0001.

